# Distinct Effects of the Extracellular Matrix on Tumor Organization and Response to Cancer Virotherapy

**DOI:** 10.64898/2026.07.20.739578

**Authors:** Ilaria Supino, Frederique Visscher, Annemarie Boerma, Anne K. Wouters, Manas Joshi, Mirjam H.M. Heemskerk, Darshak K. Bhatt

## Abstract

Extracellular matrix (ECM) regulates tumor architecture and immune accessibility, but its impact on virus-based anticancer therapies is not well understood. In particular, ECM modulation leads to both enhanced and impaired therapeutic outcomes *in vivo*, reflecting the complexity of matrix-immune-virus interactions. Here, we combine computational modeling with empirical 3D tumor spheroid-immune *in vitro* co-cultures to examine how ECM composition and abundance shape an *Alphavirus* replicon-based therapy. We show that ECM-induced changes in tumor spheroid architecture vary by cell-adhesion phenotype but higher matrix abundance consistently reduces cellular density. Notably, increasing abundance of basement membrane extract or Collagen-I reduced tumor spheroid cell density, which generally enhanced viral infection, T cell activation, and IFNγ production. However, reduced immune accessibility in collagen-rich environments suppressed immune responses despite improved infection, highlighting matrix-specific effects. Our findings provide insights into ECM’s role in virus-based therapies and propose a bottom-up framework for systematically evaluating its influence on therapeutic efficacy.

## Introduction

Virus-based therapies show promise in cancer treatment by infecting tumor cells and stimulating antitumor immune responses. However, their efficacy depends on optimal function within a tumor microenvironment shaped by extensive extracellular matrix (ECM) remodeling^1,2^. The specific ways in which ECM properties influence viral infection and immune activation remain insufficiently defined.

The ECM is an important acellular component of the tumor microenvironment, providing mechanical support while contributing to biochemical signaling that drives cancer progression^3,4^. In tumors, the ECM undergoes structural and compositional changes, including increased deposition and cross-linking of fibrillar proteins such as Collagen-I and fibronectin. These changes generate a matrix that is stiffer, denser, with marked region-to-region heterogeneity. Such variability leads to formation of micro-niches that differently influence immune evasion and tumor progression. These ECM-related features have direct implications for virus-based therapies since its efficacy depends on effective viral infection and subsequent immune activation. In addition to structural effects, ECM components and associated signaling pathways can directly influence viral infection and tumor cell behavior, including integrin-mediated viral entry and matrix-regulated signaling cascades^5,6^. Studies so far have demonstrated that a dense or highly cross-linked matrix can act as a physical barrier, limiting viral penetration into the tumor core and restricting immune cell access to infected regions^7,8^. Moreover, ECM composition can modulate antigen presentation and immune signaling, thereby influencing the magnitude and quality of antitumor immunity. For example, collagen-rich tumors have been shown to hinder T-cell migration, while altered ECM organization can reshape dendritic cell activation and cytokine production^9,10^.

Despite these observations, much of the current understanding of ECM-virotherapy interaction derives from in vivo tumor models, where ECM properties are difficult to control or isolate. Studies using oncolytic herpes simplex virus^8,11^, adenovirus^12–14^, and other viral platforms^1,7,15^ have demonstrated that matrix density and immune context influence therapeutic outcomes, yet these systems inherently couple ECM remodeling with changes in vasculature, stromal composition, and inflammatory signaling. As a result, it is challenging to understand the specific contributions of composition, abundance, or cross-linking to viral infection dynamics or immune activation. In contrast, ECM-targeting strategies, such as overexpression of matrix metalloproteinases, co-delivery of hyaluronidase, or engineering viruses to express ECM-degrading proteins, have improved viral spread in dense tumors^11–22^. However, these interventions simultaneously perturb multiple components of the TME, complicating mechanistic interpretation.

In this study, we employed an integrated bottom-up *in vitro* approach (**Fig1A-C**) combined with computational modeling (**Fig1D**) to investigate how ECM properties regulate the efficacy of virus-based therapies. This combined strategy allows controlled experimental perturbations alongside mechanistic *in silico* modeling, and is increasingly used to resolve complexity in therapeutic responses^23–30^. As a proof-of-concept, we tested the efficacy of recombinant Semliki Forest virus (rSFV)-based replicons that allow for a single round of virus infection, and encoding a fluorescent reporter to quantify viral infection kinetics and subsequent immune activation. rSFV is of particular therapeutic interest because it can be readily adapted to express immunogenic molecules^31^, and our laboratory is actively exploring rSFV-based strategies for cancer immunotherapy^25,32^. Using well-defined and human-based *in vitro* tumor-immune co-culture models^31^, we first systematically modulate ECM composition and stiffness by varying concentrations of Collagen-I and basement membrane extract, enabling controlled reconstitution of diverse matrix environments. Using this platform, we studied how matrix features influence the efficiency of both viral infection and T-cell activation. To complement mechanistic interpretation of the experimental findings, we developed a computational model simulating complex virus-tumor-immune interaction and assessed the effect of factors, such as cancer cell organization and T-cell mobility in matrix-specific microenvironments.

**Figure 1:**
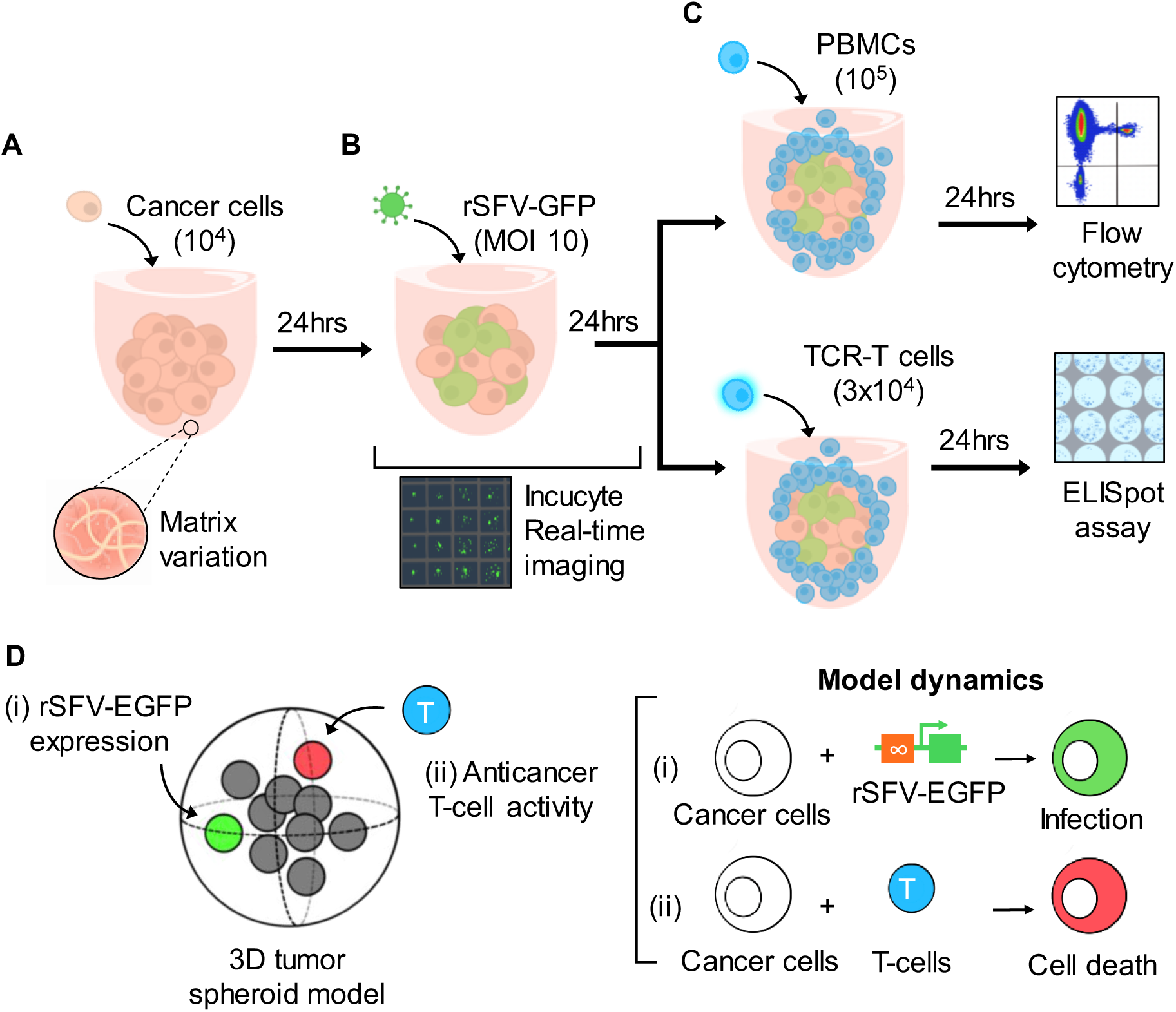
Bottom-up in vitro and in silico workflow to study the effect of extracellular matrix on cancer virotherapy. **(A)** Experimental setup to culture matrix-embedded tumor spheroids using a 96-well U-bottom plate. **(B)** Infection of tumor spheroids with rSFV-EGFP virotherapy and time-lapse microscopy to follow infection and EGFP expression kinetics over 24 hours. **(C)** Co-culture of infected tumor spheroids with PBMCs or TCR-T cells for 24 hours followed by analysis of T-cell activation. **(D)** An overview of the agent-based computational model simulating anticancer virotherapy. In a spatially explicit organization of the 3D tumor spheroid, we model two cellular dynamics (i) where cancer cells express rSFV-transgenes (e.g. EGFP or antigen) upon infection and are (ii) attacked by anticancer T-cells upon antigen expression.

## Methods

### Cell culture

22Rv1 (human prostate epithelial carcinoma), C4-2B4 (human prostate cancer), Ca-Ski (human cervical epidermoid carcinoma) and PANC-1 (human pancreatic carcinoma) cells were cultivated in RPMI complete medium (Gibco TM) supplemented with 10% (v/v) fetal bovine serum and 100 U/ml penicillin and streptomycin (Thermo Scientific). Cells were cultured in T25 flasks and passed twice a week using PBS (Thermo Fisher) for wash and 1% v/v Trypsin for cell detachment. PBMCs were thawed from liquid nitrogen in RPMI complete medium and viable cells were stained and counted using Trypan Blue. For cryopreservation, cells were frozen using fetal bovine serum supplemented with 10% DMSO (Thermo Fisher). All cultures were maintained at 37°C with 5% CO_2_.

### Analysis of gene expression

Gene expression data (calculated RNA expression) were obtained from the GSE36133 and Sanger Cell Line Project datasets. Genes are categorized into three functional groups based on their described role in cellular adhesion: cell-to-cell adhesion, cell-to-matrix adhesion, and multiple interactions. Fold-change differences between the PANC-1 and 22Rv1 cell lines in expression levels of various genes were calculated to generate the plot.

### Tumor spheroid culture in ECM

Cancer cells (22Rv1, C4-2B4, Ca-Ski, PANC-1) were counted and seeded at 10000 cells per well in a 96 well NuncSphera U-bottom plate (Corning). To generate spheroids embedded in ECM (**Fig1A**), cells were mixed with different dilutions (range 0 to10%) of either Cultrex Reduced Growth Factor Basement Membrane Extract (RGF BME, Type 2, R&D Systems, Bio-techne) or rat tail Collagen-I (Corning). Collagen-I consists of a single type of polymer, whereas BME contains a mixture of laminin, glycoprotein, Collagen-IV and heparan sulfate proteoglycan. To promote cell aggregation and tumor spheroid formation, cells were centrifuged at 1500 RPM for 10 minutes at 4°C, followed by overnight incubation.

### Design of rSFV-particles

Recombinant Semliki Forest virus (rSFV)-based replicon particles were generated using the system developed by Smerdou and Liljestrom^33^, enabling expression of enhanced green fluorescent protein (EGFP) via a single round of infection. The EGFP coding sequence (Eurofins Genomics) was cloned into the pSFV replicon backbone using PspOMI and XmaI restriction sites and propagated in E. coli JM110. Constructs were validated by Sanger sequencing (Eurofins Genomics).

### Production and titration of rSFV-EGFP viral particles

rSFV-EGFP particles were produced by linearizing pSFV-EGFP and SFV-Helper2 plasmids with SpeI (Life Technologies), followed by in vitro transcription using SP6 polymerase (Amersham Pharmacia Biotech) and a capping analogue (Life Technologies). BHK-21 cells were co-transfected with replicon and helper RNA at a 2:1 ratio by electroporation (BioRad Gene Pulser II; 2 pulses, 850 V / 25 μF). Cells were cultured in RPMI with 5% FCS and antibiotics at 30°C for 48 h. Viral particles were purified by discontinuous sucrose gradient ultracentrifugation and stored in TNE buffer at −80°C. Titration was performed by infecting BHK-21 monolayers, fixing after 24 hours with 10% acetone, staining for nsP3 (rabbit anti-nsP3, 1:2000; Cy3-anti-rabbit, 1:200), and counting positive cells by fluorescence microscopy.

### Tumor spheroid infection

Before infection, rSFV-GFP particles were activated using α-chymotrypsin (10 mg/mL, 1:20 volume; Sigma) and 2 mM CaCl₂ for 30 min, followed by inhibition with aprotinin (2 mg/mL, 1:2 volume). Spheroids were infected at MOI 10 and incubated for 24 hours (**Fig1B**).

### Tumor-immune cocultures

Infected tumor spheroids were cocultured with HLA-A*02-typed PBMCs (1E5 per well) and incubated for 24 hours. In parallel, we used CD8+ TCR-T cells, expressing MAGEA1-antigen-specific HLA-A*02:01 restricted TCR(−4F7), engineered at the Heemskerk lab at LUMC, Netherlands^34^. 3×10^4^ TCR-T cells were added per well and left in incubation for 24 hours (**Fig1C**).

### Infection analysis

Fluorescent signal of infected cancer cells (EGFP+ cells) was monitored over 24 hours of incubation using Incucyte-based time lapse microscopy. Brightfield and fluorescence images were used respectively to assess tumor spheroid area and the magnitude of EGFP expression. Cell density of each tumor spheroid was calculated by taking the ratio of number of cells and the spheroid volume.

### Quantification of T cells activation

After the coculture of HLA-matched PBMCs with infected tumor spheroids, samples were transferred to 96-well v-bottom plate (Corning) washed with 2% FBS in PBS and stained for surface activation markers. Flow cytometry was used to quantify CD4+ and CD8+ T cell activation based on CD69 and CD107a expression. Relevant antibodies are listed in **Table 1**.

**Table 1:** Antibodies used for flow cytometry.

| Antibody target | Company | Catalog # | Antibody target | Company | Catalog # |
| --- | --- | --- | --- | --- | --- |
| FITC anti-human CD4 OKT4 | Biolegend | 317408 | CXCR5-BUV563 | BD | 741316 |
| APC/Cyanine7 anti-human CD8 SK11 | Biolegend | 344714 | TIGIT-BV711 | BD | 747839 |
| APC anti-human CD69 FN50 | Biolegend | 310910 | CD24-BV480 | BD | 746278 |
| PE anti-human CD107a H4A3 | Biolegend | 328608 | CD56-BUV395 | BD | 740299 |
| Fixable Viability stain 780 | BD | 565388 | CD28-BUV496 | BD | 741168 |
| CD3-SparkBlue550 | Biolegend | 344852 | CD19-Pacblue | Biolegend | 302232 |
| CD4-cFluorYG584 | Cytek | R7-20041 | CTLA4-BV421 | Biolegend | 369606 |
| CD8-BUV805 | BD | 612889 | OX40-BV605 | BD | 745217 |
| CD45RA-Sparknir 685 | Biolegend | 304168 | CD69-BV650 | Biolegend | 310934 |
| CD95-BB700 | BD | 566542 | CCR7-BV786 | BD | 566758 |
| HLA-DR-BV570 | Biolegend | 307638 | CD137-PE | Miltenyi | 130-119-885 |
| CD38-APC-fire810 | Biolegend | 303550 | CD40L-PE-Dazzle594 | Biolegend | 310840 |
| CD27-Viobright FITC | Miltenyi | 130-113-634 | IgD-PerCP-fire-806 | Biolegend | 307818 |
| PD1-BV750 | BD | 566758 | CD160-Pe-Cy7 | Biolegend | 341212 |
| Tim3-BUV737 | BD | 748820 | ydTCR-RB744 | BD | 570495 |
| CD127-APC-R700 | BD | 565185 | TruStain Mono blocker | Biolegend | 426103 |

### Quantification of IFNγ production

ELISpot assays were performed using ethanol-activated MultiScreen HTS-IP plates (Millipore, MSIPS4510). Plates were coated overnight at 4°C with anti-human IFNγ (Mabtech, 3420-3-1000; 5 μg/mL) and blocked with X-VIVO + 2% human serum for 1 h at 37°C. Cells from TCR-T cocultures were plated and incubated for 24 hours. Spots were developed using biotinylated anti-IFNγ (Mabtech, 3420-6-1000; 1 µg/mL), streptavidin-poly-HRP (Sanquin, M2051), and TMB substrate (Mabtech, 2651-10).

### Flow cytometry based deep T cell phenotyping

Spectral flow cytometry was performed to characterize T cell populations using a multi-parameter antibody panel targeting surface markers. Single-cell data were exported from cytometry acquisition software as CSV files and analyzed in R (v4.3.0). Individual sample files were imported and merged into a single dataset, with sample identifiers retained. Non-informative parameters (e.g., time, autofluorescence, event metadata) were excluded, and marker expression values were retained for downstream analysis. To reduce computational burden while preserving population structure, a random subset of 25% of total events was selected. Marker expression values were z-score scaled, followed by dimensionality reduction using principal component analysis (PCA). The top principal components (up to 30) were used as input for Uniform Manifold Approximation and Projection (UMAP) to generate a two-dimensional embedding of single-cell data. Unsupervised clustering of UMAP coordinates was performed using k-means clustering (k = 5) to define phenotypically distinct T cell populations. Cluster frequencies were calculated per experimental condition and expressed as fractions of total cells. Marker expression patterns were analyzed at single-cell resolution by transforming the dataset into long format and calculating z-score–normalized expression per marker. Mean marker expression per cluster and condition was computed and visualized using heatmaps. Additionally, population distributions and marker expression patterns were visualized on UMAP embeddings using density plots and binned expression maps.

### Computational model overview

We developed a stochastic agent-based model, called sa-mRNA Immunotherapy Model (SAMI), implemented in R (v4.3.0)^35^ to simulate the interaction between rSFV-viral infection (modeled as self-amplifying sa-mRNA expression), cancer cells and T cell responses (See **Fig1D** for an illustrated overview of the model). In particular, we modelled viral gene (e.g. EGFP or an antigen) expression in tumor spheroids and activation of cytotoxic T cells leading to tumor-killing. Each simulation tracked EGFP expression at a single-cell resolution, its modulation by microenvironmental factors, and T cell-mediated cytotoxicity. Simulations were run with hourly timesteps covering a 24 hours window to capture EGFP kinetics and immune interactions over time. Please refer to **Supplementary table 1** for model parameters and details of the simulations.

Tumor spheroids were generated by uniformly seeding 10,000 cells in a sphere with a range of 50 to 500 spheroid radius (*R*) cell units, allowing us to systematically modulate global tumor organization and density ranging from 10^−3^ to 10^−6^ cells per spheroid. ECM-driven changes in tissue architecture were captured through variations in both spheroid size and local cellular crowding. For each cell (*i)*, spatial coordinates *x_i_* = (*x_i_*, *y_i_*, *z_i_*) were sampled such that ||*x_i_*|| ≤ *R*. Local crowding was quantified as the weighted density of neighboring cells within a 10-unit radius using a Gaussian-weighted crowding density, defined as 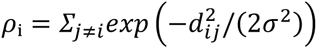, with *σ* = *r_c_* / 2, where *d_ij_* = ‖*x*_i_ − *x*_j_‖ and *r_c_* denotes the crowding radius. This density was converted into a crowding factor (*C_i_*) that inhibited EGFP expression, such that cells in densely packed regions exhibited reduced expression compared with those at the periphery. Here, *C*_i_ = 1/(1 + *β*(*ρ*_i_/*ρ_ref_*)), and *ρ_ref_* is the median crowding across cells and β controls the strength of ECM-induced inhibition.

Viral infection was initiated by assigning each cell a stochastic input of sa-mRNA, modeled as a Poisson process with effective dose *D*_i_ = *D* · *f_delivery_*(*x*_i_) · *C*_i_, such that infection probability depends on both spatial delivery and local crowding. Subsequent intracellular dynamics were governed by two principal kinetic parameters: the genomic replication rate k₁ and the subgenomic transcription rate k₂. These processes were modeled as *ΔR*_i_(*t*)∼*Poisson*(*k*_1_ · *m*_i_), *Δs*_i_(*t*)∼*Poisson*(*k*_2_ · *R*_i_), linking viral replication to downstream protein production. EGFP accumulation followed saturating kinetics as *ΔP*_i_(*t*)∼*Poisson*(*s*_i_ · (1 − *P*_i_/*P*_i_*max*) · *γ*_i_ · *I*_i_ · *C*_i_). Here *P*_i_, *max* represents the maximum protein capacity of a cell and γ_i_ captures intrinsic variability in translational efficiency.

To evaluate how ECM-driven organization affects immune responses, T cell-mediated cytotoxicity was incorporated explicitly. T cells were seeded randomly within the spheroid and could only interact with target cells within a finite spatial radius defined by *d_ik_* ≤ *p_kill_*_,*i*_ making spatial proximity a critical determinant of immune activity. Killing probability depended on intracellular antigen levels relative to a stochastic recognition threshold, *θ*_i_∼*Uniform*(*θ*(1 − *ε*), *θ*(1 + *ε*)), and was further modulated by microenvironmental constraints such that *p_kill_*_,*i*_ ∝ *P*_i_ · *C*_i_. Thus, regions of high crowding not only suppress viral expression but also reduce effective T-cell killing, linking ECM structure to immune regulation.

Each T cell was limited to a maximum number of killing events (*K_max_* = 5), representing finite cytotoxic capacity^36,37^. This constraint becomes particularly important in densely packed tumors, where restricted access (low *p_kill_* effectiveness due to crowding) and reduced antigen expression jointly limit immune-mediated clearance.

A total of 100 replicate spheroids were simulated to account for stochastic variability. Outputs included single-cell EGFP expression levels, spatial distributions of expression, and cell-survival status at each time point. Effects of cellular–crowding and ischemia on EGFP expression were quantified using LOESS regression. T cell activity was measured as the number of kills per T cell, and overall tumor cytotoxicity was calculated as the fraction of cells killed at endpoint (at 24 hours), stratified by density of neighboring cancer cells. Data compilation and visualization were performed using the R packages ggplot2 and patchwork.

### Statistical analysis

Incucyte image data were analyzed using the IncuCyte Live-Cell Analysis software (Sartorius). Cell density values were calculated and log₁₀-transformed prior to analysis. Flow cytometry data were processed in FlowJo (FlowJo LLC). All plots and statistical analyses were generated in GraphPad Prism 9 (GraphPad Software) and RStudio. Bivariate linear regression with 95% confidence intervals was used to assess correlations, with correlation coefficients (r) and p-values reported in each plot. Comparisons between two independent groups were performed using two-tailed unpaired t-tests, while multiple-group comparisons were evaluated using one-way ANOVA. Statistical significance was defined as p < 0.05.

### Data and code availability

The SAMI model code for a terminal based version and an executable SHINY version are available on www.github.com/d-bhatt/SAMI.

## Results

### Modeling cancer cell organization in the presence of extracellular matrix

We studied how the presence, abundance and type of extracellular matrix (ECM) changes organization of cancer cells in three dimensional space using spheroid-based culture system **(Fig2A)**. Cultivation of 22Rv1 cancer cells in basement membrane extract (BME) or Collagen-I matrix led to decrease in spheroid density with increasing matrix abundance **(Fig2B-C)**. The density remained unchanged over the observed time-period of 24 hours **(Fig2D)**. Of note, ECM-driven reduction in cell density was more pronounced in BME-embedded cultures compared to those in Collagen-I. Cells in BME formed sparsely organized clumps, whereas Collagen-I supported the formation of a larger, though less dense spheroid structure (**Fig2B**).

**Figure 2:**
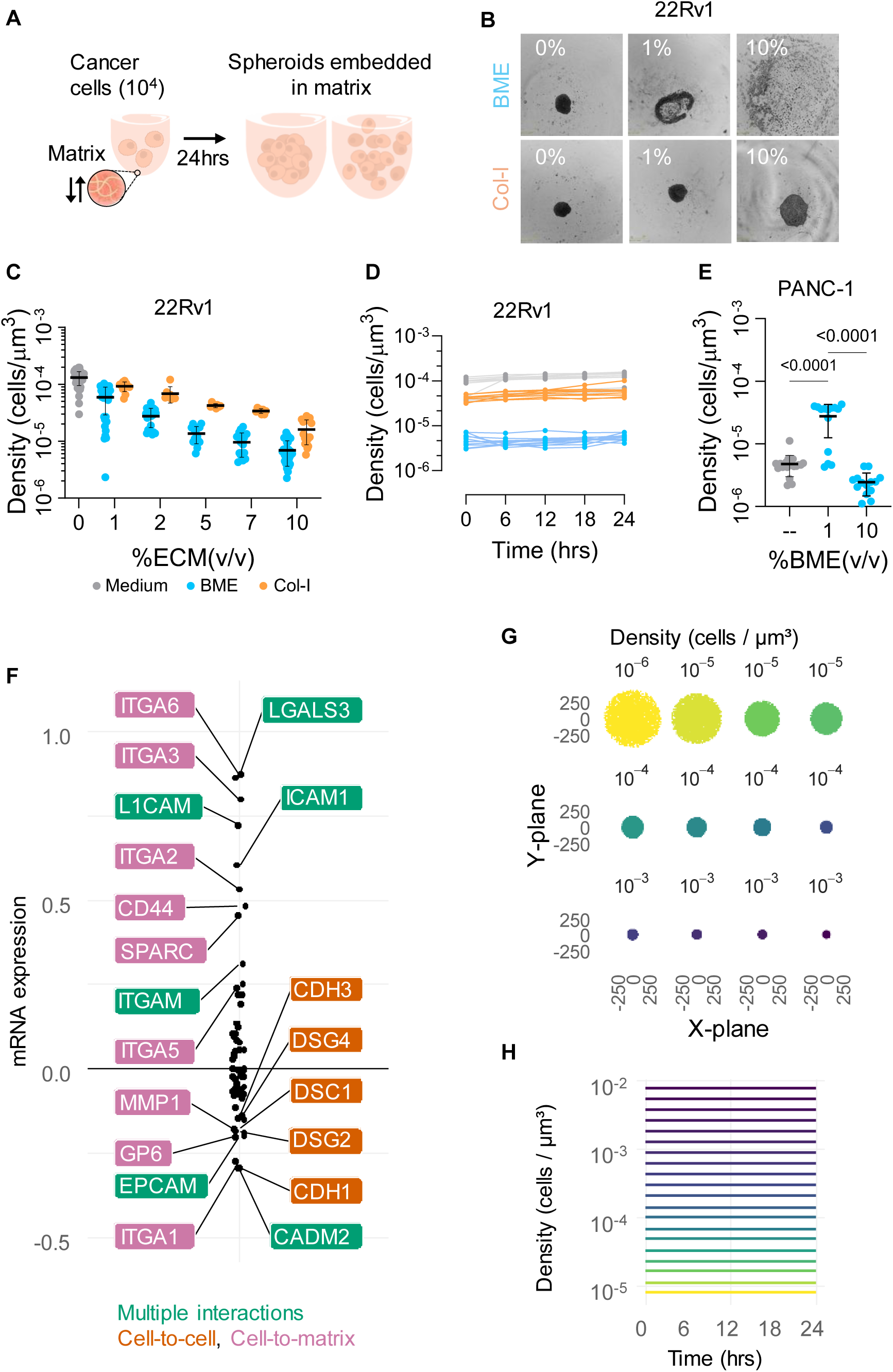
Effect of extracellular matrix on cancer cell organization and density. (**A**) Schematic experimental setup to culture tumor spheroids in different matrices over 24 h, resulting in spheroids of different cell densities. (**B**) Live-cell imaging of 22Rv1 tumor spheroids embedded in 0%, 1% and 10% BME or Collagen-I. (**C**) Effect of BME or Collagen-I embedding (matrix abundance 0-10%) on 22Rv1 spheroid cell density. (**D**) Kinetics of spheroid cell density over 24 hours of culture. (**E**) Effect of BME embedding on PANC-1 spheroids cell density. (**F**) Gene expression profile comparison between PANC-1 and 22Rv1 cancer cell lines using data from CellExpress database^38^. Genes are colored by interaction type: multiple interactions (green), cell-cell interaction (orange), cell-matrix interaction (purple). (**G**) Spheroids of different densities simulated in silico using a computational model. (**H**) Kinetics of spheroid density over 24 hours in simulated spheroid models. Replicate number varied by condition across plots; all conditions had n>8, and some conditions used n>15. In (**E**) statistical analysis was performed using ordinary one-way ANOVA followed by multiple comparisons using the two-stage linear step-up procedure of Benjamini, Krieger and Yekutieli. Statistical significance was defined as p < 0.05.

A cell-type dependent effect was observed at lower matrix abundance (1%), with an increase in cancer cell density for PANC-1 and Ca-Ski tumor spheroids (**Fig2E** and **Supplementary FigS1**). However, higher matrix abundance (10%) generally decreases density across multiple cancer cell lines (**Fig2E** and **Supplementary FigS1**). To explore why 22RV1 and PANC-1 cells display distinct cell-organization and density patterns, we assessed differences in adhesion-related gene expression patterns. Expression of adhesion molecules differed between cell types at baseline, where PANC-1 cells expressed more cell-to-matrix adhesion related molecules, whereas 22Rv1 expressed more cell-to-cell adhesion related molecules **(Fig2F)**. Using these insights, we employed an agent-based modeling approach to simulate virtual tumor spheroids of different densities **(Fig2G)** maintaining their density over time **(Fig2H)**.

### Modeling cancer cell infection in the presence of extracellular matrix

Using rSFV-EGFP as a model virotherapy **(Fig3A)**, we tested the effect of ECM-mediated changes in cell organization on viral infection in the spheroid-based culture system. Cancer cells embedded in different matrix abundances were infected and EGFP expression per cell was quantified using Incucyte-based real-time microscopy imaging (**Fig3B**). An increase in the frequency of EGFP+ cells was observed in 22Rv1 tumor spheroids with lower density (**Fig3C-D**). Presence of either BME or Collagen-I matrix led to an increase in viral infection compared to spheroids without matrix-embedding **(Fig3D)**.

**Figure 3:**
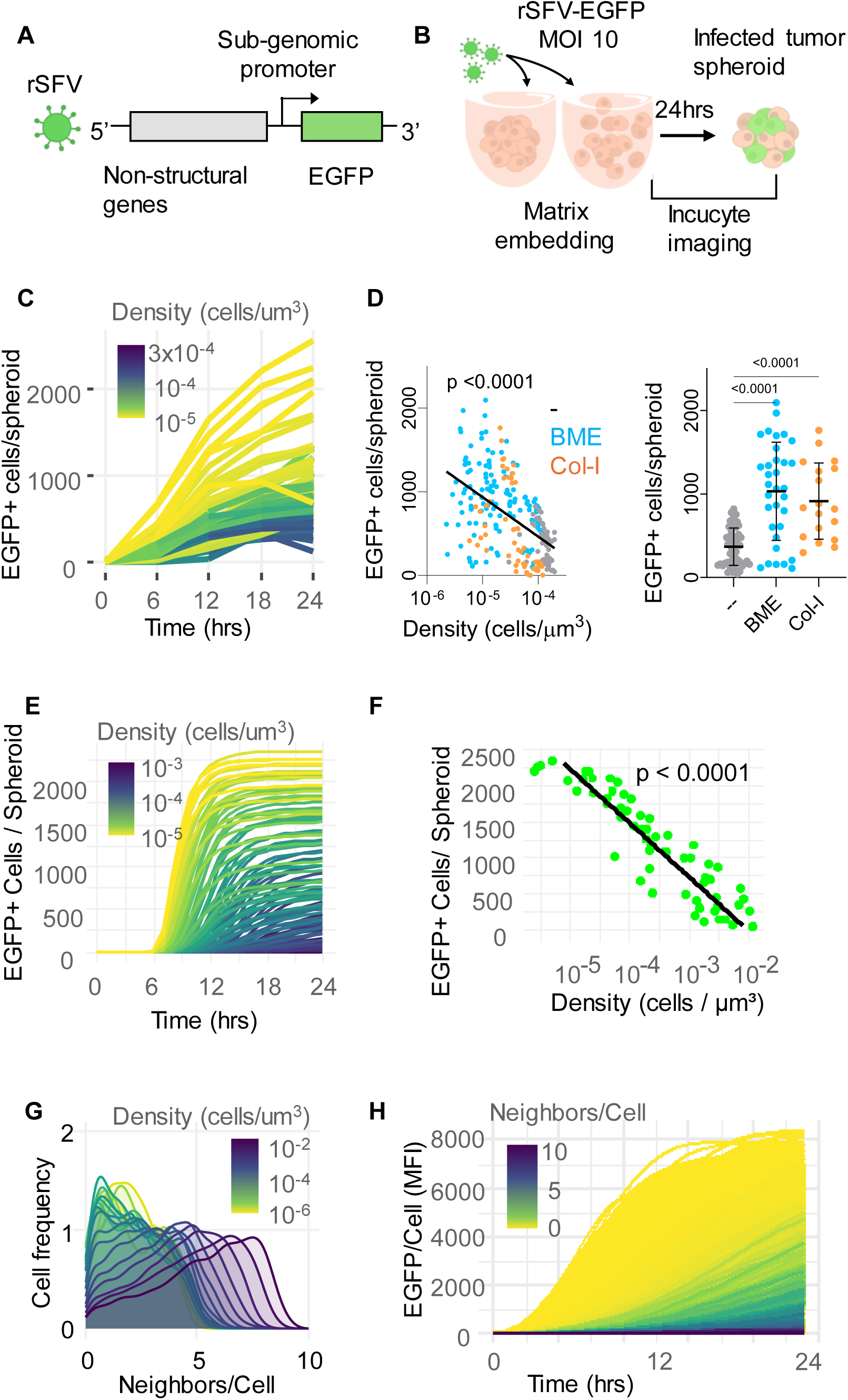
Effect of cancer cell-density and extracellular matrix on virotherapy infection kinetics. (**A**) Illustration of rSFV replicon encoding EGFP and capable of a single round of infection. (**B**) Schematic experimental setup to culture tumor spheroids embedded in matrices followed by infection with rSFV-EGFP virus particles. (**C**) Number of EGFP positive cells per spheroid over 24 hours of infection, where spheroid density is indicated by a color gradient. (**D**) Correlation between rSFV-infection and spheroids density along with a comparison of spheroids with or without matrix embedding. Figures **E-F** show results from computational model simulation. (**E**) Number of EGFP positive cells per spheroid over 24 hours of infection, where spheroid density is indicated by a color gradient. (**F**) Correlation between rSFV-infection and spheroid density after 24 hours of simulation. (**G**) Neighboring-cell frequency distribution across spheroids of different cell densities. (**I**) Level of EGFP expression per cell with color gradient showing number of neighbor cells. Replicate number varied by condition across plots; all conditions had n>8, and some conditions used n>15. In (**D**) statistical analysis was performed using linear regression. Plot (**E**) was analyzed using ordinary one-way ANOVA followed by multiple comparisons using the two-stage linear step-up procedure of Benjamini, Krieger and Yekutieli. Statistical significance was defined as p < 0.05.

A similar trend of increasing infection with lower cell density was observed in simulations using the *in silico* model **(Fig3E-F),** when considering crowding radius of 3-5 cell units (**Supplementary FigS2**). At a single-cell resolution *in silico*, we observed that cells with more neighbors were frequently found in dense spheroids **(Fig3G)** which in turn correlated to a reduction in EGFP expression **(Fig3H)**. Of note, EGFP levels per cell were highly heterogeneous, shaped by spatial crowding and ischemia-like gradients, where larger tumor spheroids and regions with higher local neighbor density accumulated EGFP more rapidly over time. Spatial visualization at 24 hours revealed that peripheral cells consistently reached higher EGFP levels than core cells (see **Supplementary FigS2** for representative images).

### Quantifying T-cell activation in the presence of extracellular matrix

We cocultured rSFV-EGFP-infected cancer cells embedded in BME and Collagen-I matrices with HLA-matched PBMCs for 24 hours and quantified T cell activation through flow cytometry **(Fig4A)**. Activation was defined as expression of both CD69 and CD107a by helper (CD4+) and cytotoxic (CD8+) T cells. CD4+T cell activation did not correlate with changes in cell density, whereas CD8+T cell activation increased significantly as density decreased **(Fig4B)**. Cocultures in presence of 10% BME showed higher CD4+ and CD8+ T cell activation than non-embedded controls. In contrast, embedding in 10% Collagen-I produced fewer or no changes in CD4+ T cell-activation and a significant reduction in CD8+ T cell-activation **(Fig4C)**. These results were observed across different target tumor cells **(Supplementary FigS3)** and were independent of the rSFV-infection or transgene expression **(Fig4D** and **Supplementary FigS3)**, suggesting direct ECM-mediated effects.

**Figure 4:**
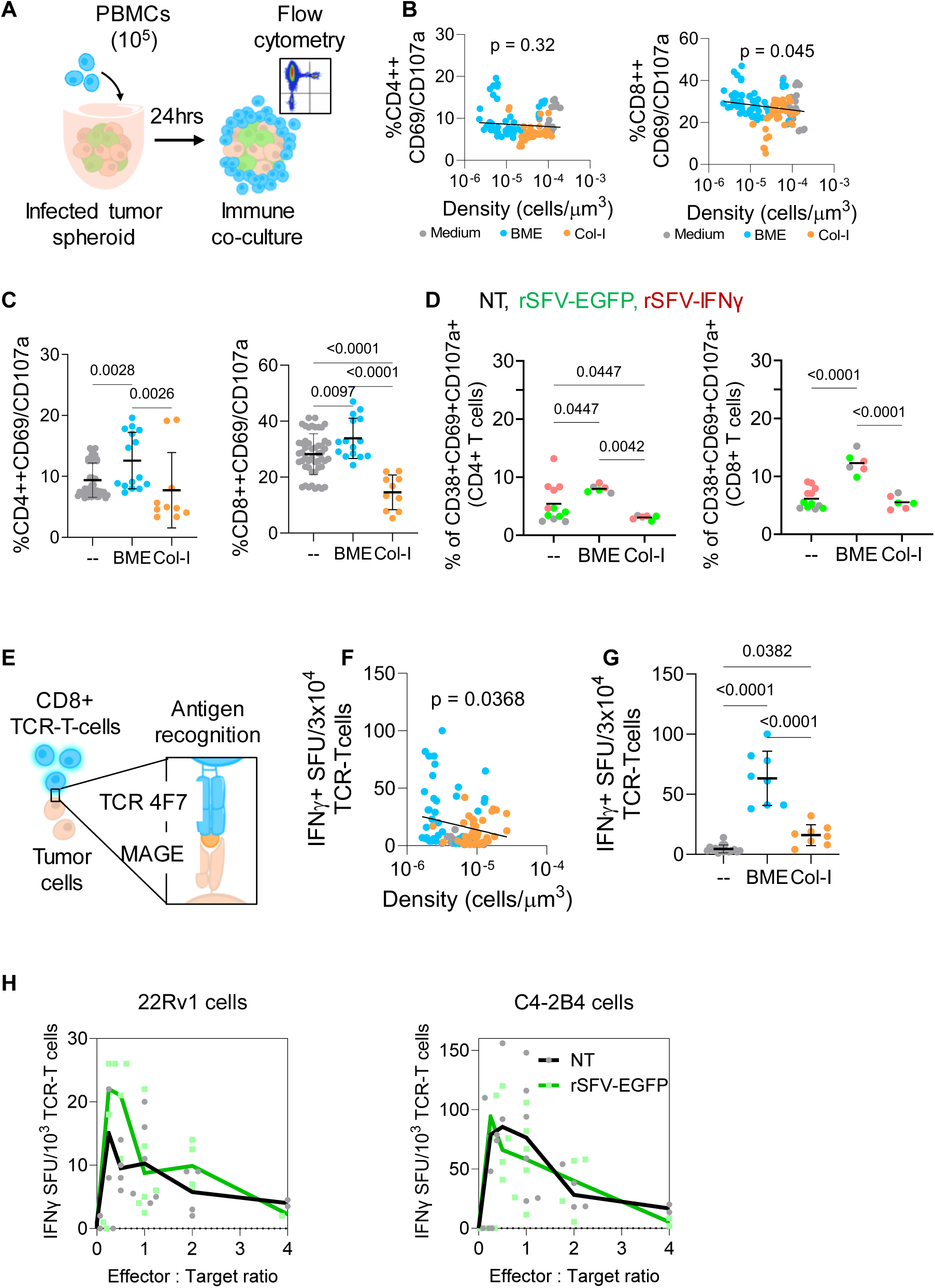
Effect of cell density and extracellular matrix on T cells activation and cytokine secretion. (**A**) Schematic of the experimental setup to co-culture rSFV-infected spheroids with HLA-matched PBMCs and Flow cytometry analysis. (**B**) Correlation of 22Rv1 tumor spheroids cell density with percentage of CD4+ or CD8+ T cells expressing CD69 and CD107a. (**C**) Comparison of CD4+ and CD8+ T cell activation upon co-culture with 22Rv1 spheroids in presence of BME or Collagen-I matrix. (**D**) Percentage of CD4+ or CD8+ T cells expressing CD38, CD69 and CD107a upon co-culture with 22Rv1 spheroids in presence of BME and Collagen-I matrix. Colors indicate spheroid status: uninfected (NT, grey), rSFV-EGFP-infected (green), or rSFV-IFNγ-infected (red). (**E**) Schematic of the experimental co-culture of tumor spheroids with CD8⁺ TCR-T cells recognizing the MAGEA1 antigen expressed by 22Rv1 cells. (**F**) ELISpot assay performed 24 h after co-culture, showing IFNγ spot-forming units (SFUs) of T cells as a function of 22Rv1 tumor spheroid cell density. (**G**) Comparison of IFNγ SFUs of T cells co-cultured with 22Rv1 spheroids without matrix or embedded in BME or Collagen I. (**H**) IFNγ SFUs of T cells co-cultured with 22Rv1 and C4-2B4 tumor spheroids at increasing effector-to-target ratios, comparing uninfected spheroids with those infected with rSFV-EGFP. Replicate number varied by condition across plots; all conditions had n>6, and some conditions used n>10. Plots (**B** and **F**) were analyzed using linear regression. Plots (**C**, **D** and **G**) were analyzed using ordinary one way ANOVA followed by multiple comparisons using the two-stage linear step-up procedure of Benjamini, Krieger and Yekutieli. Statistical significance was defined as p < 0.05.

To study antigen-specific T cell responses, we cocultured infected ECM-embedded cancer cells expressing MAGE antigen along with CD8+ T cells expressing the corresponding cognate-TCR-4F7 receptor **(Fig4E)**. IFNγ spot-forming units, as a measure of T cell activation, increased as cell density decreased **(Fig4F)**. Embedding in 10% BME enhanced IFNγ secretion relative to non-embedded cocultures, whereas embedding in 10% Collagen-I markedly reduced cytokine production compared to BME-embedded samples **(Fig4G)**. Again, these effects were independent of rSFV-infection and were observed across different target tumor cells **(Fig4H)**.

### Changes in overall T-cell phenotype in the presence of extracellular matrix

To understand why BME and Collagen-I differentially modulate T-cell activation, we analyzed T-cell phenotypes after coculturing PBMCs with tumor spheroids embedded in each matrix type. Using a panel of surface markers (**Supplementary FigS4,5**), we identified five distinct T-cell populations across embedded and non-embedded conditions **(Fig 5A)**. Supp FigS3 illustrates the expression of each surface marker at a single cell resolution. Matrix embedding, regardless of type, altered markers associated with motility (CCR7, CXCR5) and activation (OX40, CD24, CD107a, CD38, CD95). However, the two matrices produced divergent effects on exhaustion and differentiation. Collagen-I abundance correlated with increased expression of exhaustion markers (CTLA4) and terminal differentiation markers (CD45RA, CD160), whereas BME embedding was associated with reduced expression of exhaustion markers (TIGIT, PD1) **(Fig 5B)**. Furthermore, these observations were independent of rSFV-infection of the spheroids **(Supplementary FigS4)**. These differences suggest that matrix composition, not just matrix presence, plays a key role in shaping T-cell functional states.

**Figure 5:**
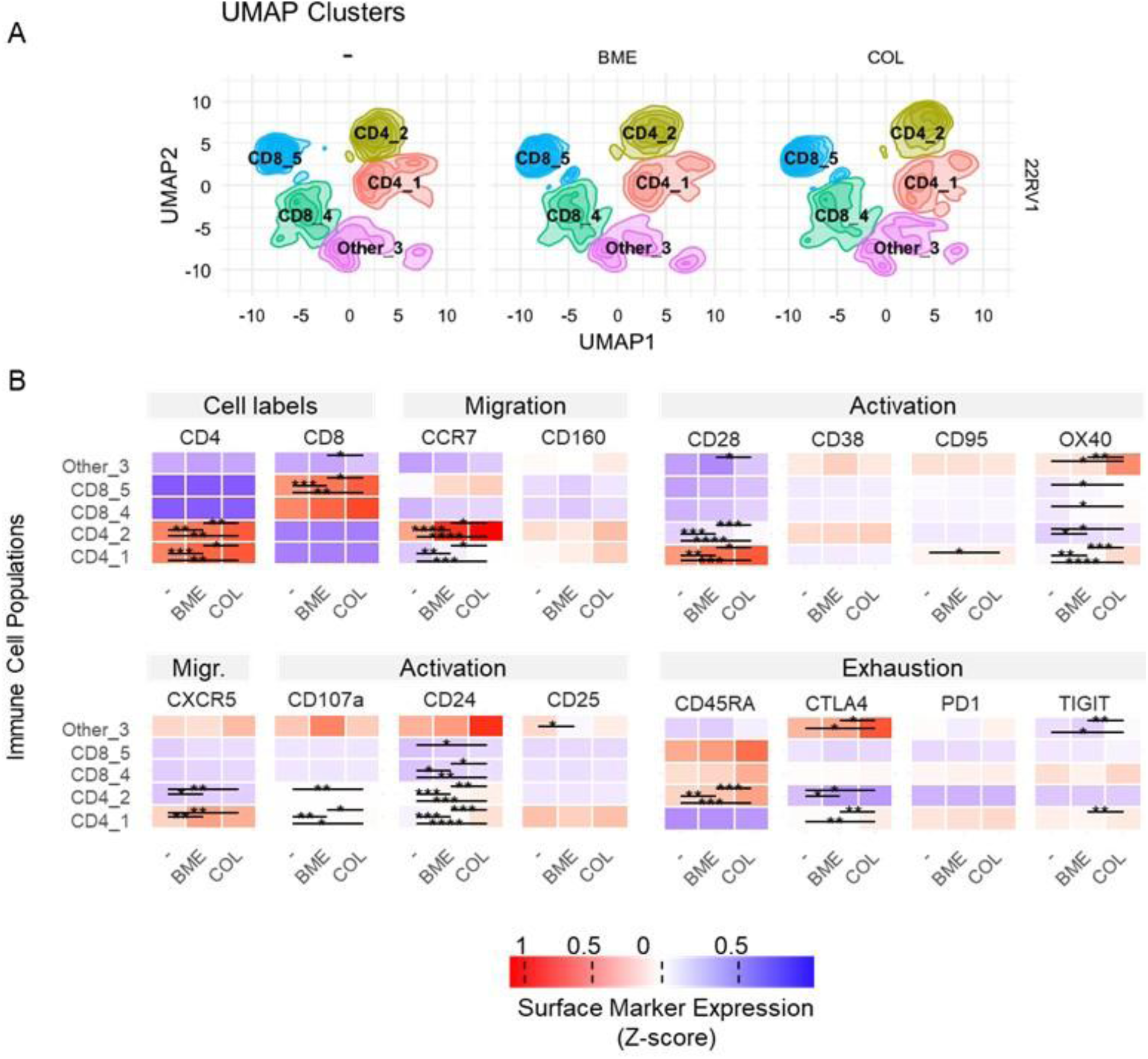
Changes in overall T-cell phenotype in the presence of extracellular matrix. Co-culture of rSFV-infected 22Rv1 tumor spheroids with HLA-matched PBMCs for 24 hours. (**A**) Comparison of UMAP cluster populations of CD4+ and CD8+ T cells between samples without matrix embedding, with BME matrix embedding and with Collagen-I matrix embedding. (**B**) Normalized expression (MFI, z-score) of selected markers involved in T cell migration, activation, and exhaustion. Statistical significance was defined as p < 0.05.

### Agent-based modelling of the effect of cell organization on T-cell responses

To evaluate how ECM-driven changes in tumor cell organization influences T-cell activation, and to understand why T-cell activation shows weaker correlations with cell-density than infection, we extended our agent-based modelling framework to incorporate T-cells. Here, we simulated T-cell infiltration and killing to determine how spatial constraints modulate antitumor immune efficacy. In particular, we studied the effect of how T-cell migration and killing across different distances influences T-cell activation and cancer cell survival in the model. As illustrated in **Fig6A**, a shorter T-cell killing radius (*K_r_*), e.g. *K_r_*=10 would lead to killing of 3 target cells, whereas a longer radius, e.g. *K_r_*=50 would lead to killing of 7 target cells.

**Figure 6:**
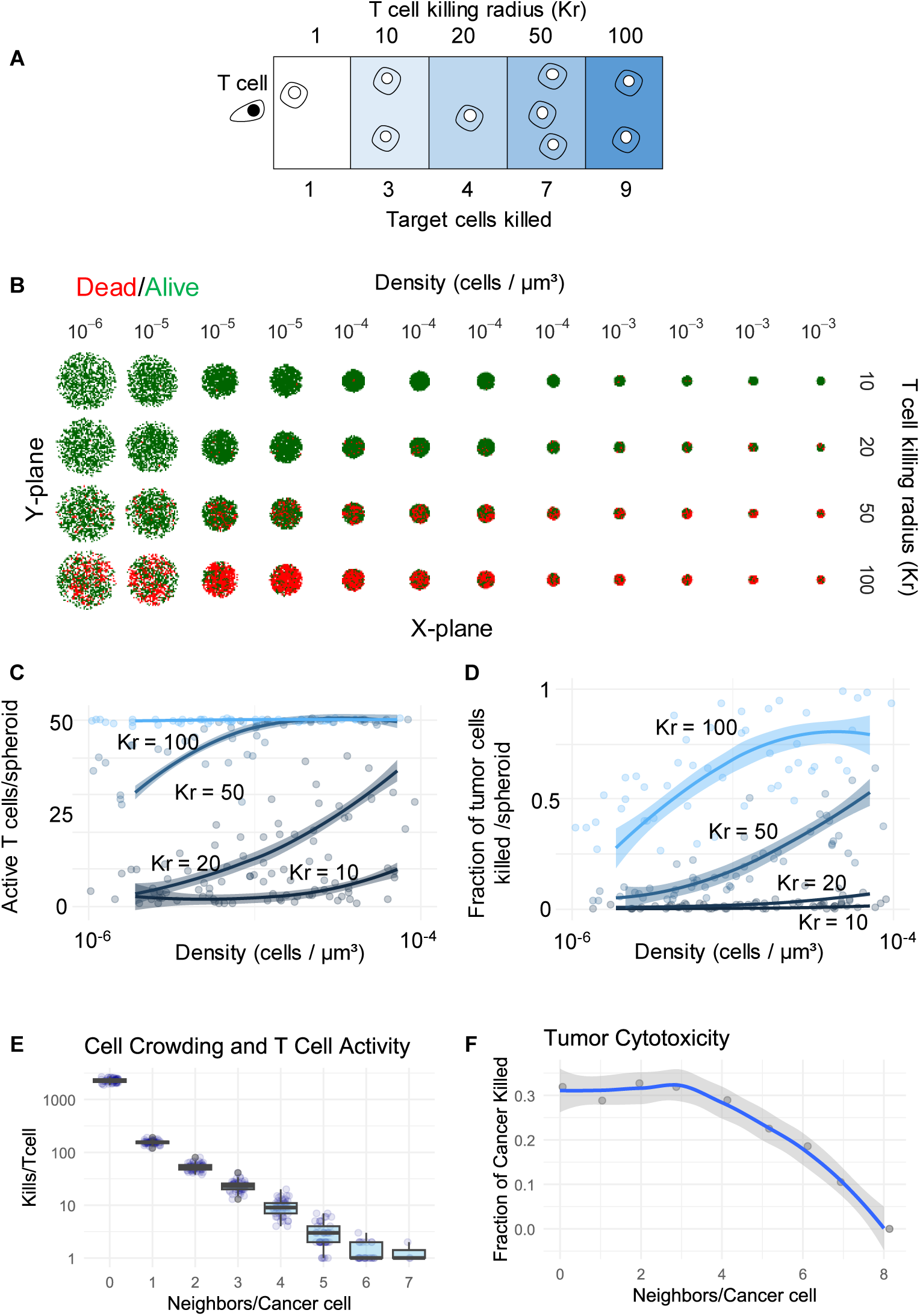
Computational modeling of the effect of cell density on anticancer T cell response. (**A**) Schematic representation of number of target cancer cells killed per T cell at different killing radius (K_r_). (**B**) Simulation of cancer cells death per spheroids of varied density and T cell killing radius. (**C**) Frequency of activated T cells per spheroid with increasing spheroid density and T cell killing radius. (**D**) Fraction of cancer cells killed per tumor spheroid with increasing density and T cell killing radius. Effect of the frequency of neighboring cancer cells on the (**E**) number of cancer cells killed per T cell and (**F**) total fraction of cancer cells killed.

Across tumor spheroids of different radii, cancer cell death was distributed heterogeneously, with killing occurring more often at the periphery in larger, less dense spheroids **(Fig6B)**. The frequency of activated T cells per spheroid increased with increasing killing radius and increasing local neighbor density **(Fig6C)**, indicating that densely packed regions provide more accessible targets for serial killing. Similarly, increasing the T-cell killing radius led to efficient killing across tumor spheroids of different densities **(Fig6D)**. However, at a single cell resolution, we observed that the killing efficacy of a T-cell decreased with an increasing number of neighboring cancer cells **(Fig6E)** correlating with a reduction in overall tumor cell killing **(Fig6F)**.

Together, these simulations suggest that the effect of cancer cell organization on T-cell activation is partially buffered by spatial limitations on T-cell access and killing, explaining the more modest experimental effects observed.

## Discussion

In this study, we set out to understand how the extracellular matrix shapes the efficacy of anticancer virotherapy by regulating cancer cell organization, viral infection, and T-cell activation. ECM remodeling is a hallmark of tumor progression and has complex effects on therapeutic resistance and efficay^3,4^. While the influence of ECM-degradation during cancer virotherapy is explored^39–41^, its exact influence in viral infection and immune activation remain less defined. By combining a controlled 3D *in vitro* system with computational modelling, we show how matrix composition and abundance modulate the spatial arrangement of cancer cells and, in turn, the dynamics of viral infection and immune activation.

Our results demonstrate that ECM abundance reorganizes tumor spheroids, thereby directly influencing infection kinetics. Increasing concentrations of BME or Collagen-I reduced cell density, but the magnitude and direction of this effect depended on both matrix type and the intrinsic adhesion phenotype of the cancer cells. BME, a softer and less structured matrix, induced pronounced cell-dispersion, whereas the stiffer and more organized Collagen-I produced more compact tumor spheroid structures. These observations are consistent with prior work showing that ECM stiffness and composition regulate tumor architecture and cell organization^4,42^. Simultaneously, transcriptomic analysis revealed that embedding different cancer cells in the same matrix can lead to distinct cell organization depending on the distinct adhesion phenotypes: PANC-1 cells, which express high levels of cell-to-matrix adhesion genes, compact at low BME levels before dispersing, while 22Rv1 cells, which rely more on cell-to-cell adhesion, disperse more uniformly. These findings highlight that ECM abundance and cancer cell adhesion jointly determine tumor architecture.

Cell reorganization had a direct impact on viral infection. Lower-density spheroids exhibited higher infection frequencies across cancer cell types. Our computational model recapitulated these dynamics, showing that reduced neighboring cell density led to earlier and higher EGFP expression at both a cellular and spheroid level. These insights help explain why infection correlated strongly with cell organization across multiple cancer cell lines. Notably, the effect of ECM on infection was matrix-independent: both BME and Collagen-I enhanced infection across several lines, corresponding to a lower cell density. Our findings are consistent with prior work demonstrating that ECM remodeling, for example via MMP activity, enhances virotherapy efficacy by improving intra-tumoral spread and accessibility^17,18,21,22^. However, our results further suggest that ECM-mediated effects extend beyond simple barrier removal^8,43^ and include indirect regulation through tumor architecture and cellular organization. This refines the prevailing view that ECM uniformly hinders virotherapy, instead revealing that the cellular organization and local cell-density determine whether it supports or restricts viral infection. Notably, our work may explain why some studies observe that inhibiting MMP activity does not necessarily impair, and may even enhance, therapeutic outcomes, highlighting the complex and context-dependent role of ECM remodeling in cancer virotherapy^44–46^.

T-cell activation followed similar but less pronounced trends with respect to cell density. In 22Rv1 spheroids, lower cell density and BME embedding enhanced CD8+T cell activation and IFNγ secretion, whereas Collagen-I reduced both responses, even below levels observed without matrix. These differences likely reflect the impact of environmental stiffness on T-cell motility and activation, as rigid matrices, like Collagen-I, can restrict immune cell infiltration and proliferation^9,47,48^. However, the weaker correlation between cell density and T-cell activation compared to infection suggested that additional constraints shape immune responses. In depth T-cell phenotyping supported this interpretation, while matrix presence increased markers of motility and activation, exhaustion and differentiation signatures diverged sharply by matrix type. Notably, Collagen-I but not BME, increased exhaustion marker expression as noted before^10^, indicating that ECM composition modulates T-cell functional states independently of infection levels. Our findings are further supported by real-time imaging studies that demonstrate how tumor microenvironment composition directly regulates T-cell migration patterns and infiltration efficiency^49^, while matrix stiffness and dimensionality further modulate T-cell activation states^9,48^.

Finally, our computational modelling provided mechanistic insight into these observations. While viral infection was highly sensitive to spatial organization, T-cell killing was constrained by physical access, for example due to a stiffer matrix, and largely saturated at the spheroid periphery. Killing efficiency decreased with spheroid radius and local neighbor density but increased with T-cell killing radius. These simulations suggest that T-cell responses are limited more by spatial accessibility than by antigen availability, explaining why T-cell activation effects were less pronounced and more variable across cell lines. This is consistent with previous observations that population context and spatial organization strongly influence infection dynamics at the single-cell level^24^. Together, the modelling and experimental data reveal that ECM-driven changes in cancer cell density predominantly influence viral infection, whereas ECM-mediated alterations in matrix stiffness and tissue accessibility primarily regulate T-cell cytotoxicity. Although prior computational models have explored ECM–virus interactions^23,50^, our work uniquely integrates these effects with immune dynamics at single-cell resolution.

To conclude, our results demonstrate that ECM composition and abundance regulate cancer virotherapy through combined effects on cancer cell organization, viral infection, and T-cell activation. By integrating a bottom-up experimental and computational framework, we provide novel insights into how viral infection is governed primarily by matrix-induced changes in cell organization, whereas T-cell activation is directly controlled by matrix composition and stiffness. This mechanistic separation highlights opportunities to independently optimize infection and immune engagement in future anticancer virotherapy designs.

## Limitations

Our bottom-up experimental *in vitro* system enabled precise control over ECM abundance and composition, allowing us to isolate their effects on cell organization, infection, and immune activation. However, this reductionist approach excludes many regulatory factors present *in vivo*, including stromal interactions, vascularization, and complex immune networks. In addition, because we employed rSFV replicons that support only a single round of infection, our study does not capture how ECM or cell organization might affect multicycle viral propagation, secondary infection waves, or long-term viral–immune dynamics. As a result, some observations may differ from *in vivo* behavior. Computational modeling helped contextualize these findings, but further integration with more complex systems will be required to fully capture ECM-immune dynamics across multiple infection cycles. Despite these limitations, the experimental control afforded by our *in vitro* model design was essential for disentangling the individual contributions of ECM composition, cell organization, and immune accessibility, providing insights that would be difficult to resolve in more complex, heterogeneous *in vivo* settings.

## Acknowledgements

This study was partially funded by ERASMUS mobility grant awarded to IS and by Dutch Council of Research (NWO) (grant# OCENW.XS25.1.147) awarded to DKB. Part of the work has been performed at the UMCG Imaging and Microscopy Center (UMIC) and the Flow Cytometry Research Unit (FCU) which is sponsored by UMCG. We thank Thijs Janzen (University of Groningen) for reviewing the code of the SAMI computational model and providing critical feedback, and Debbie van Baarle (UMCG) for helpful discussions on T cell responses.

## Author contributions

All authors made substantial contributions to the manuscript.

IS: Conceptualization, Writing and editing manuscript, Data collection and analysis, Validation, Visualization

FV: Editing manuscript, Data collection and analysis

AB: Editing manuscript, Data collection and analysis

AKW: Data collection and analysis, Validation

MJ: Editing manuscript, Software code review, Data collection and analysis, Visualization

MJMH: Conceptualization, Writing and editing manuscript, Supervision

DB: Conceptualization, Writing and editing manuscript, Data collection and analysis, Software, Visualization, Supervision, Funding acquisition

## Resource availability

### Lead contact

Further information and requests for resources and reagents should be directed to and will be fulfilled by the lead contact Darshak K. Bhatt

### Material availability

All unique/stable reagents generated in the study are available upon request, except where restricted by institutional agreements or MTAs.

### Data and code availability

All data reported in this paper will be shared by the lead contact upon request. Any additional information required to reanalyze the data reported in this paper is available from the lead contact upon request. The code used for this work and an executable version of the SAMI model can be found at https://github.com/d-bhatt/SAMI.

