## Supplementary table and figures for "Distinct Effects of the Extracellular Matrix on Tumor Organization and Response to Cancer Virotherapy"

**Supplementary table 1**

| Parameter | Interpretation | Default value | Evaluated range |
|---|---|---|---|
| $t$ | Simulation duration representing the time window for sa-mRNA expression dynamics and immune interactions | 24 h * | 0–24 h |
| $N\_cells$ | Number of cancer cells forming the tumor spheroid | 10000 * | 1000-10000 |
| $R$ | Tumor spheroid radius controlling global tumor size and cell density | - | 50–500 cell units* |
| $r_c$ | Crowding radius defining the local neighborhood contributing to ECM/cell-density effects | 5 | 1–10 cell units |
| $C_i$ | Crowding factor representing inhibition of antigen expression and T-cell activity caused by local cell density | - | 0 or 1 |
| $B$ | Strength of ECM/crowding-mediated inhibition | 1 | - |
| $D$ | Initial sa-mRNA therapeutic dose delivered to tumor cells (corresponding to Multiplicity of infection: MFI) | 10 * | - |
| $k_1$ | Genomic replication rate of self-amplifying sa-mRNA | 0.5 * | - |
| $k_2$ | Subgenomic transcription rate controlling antigen production | 1.0 * | - |
| $p_{kill}$ | Maximum distance allowing T-cell interaction and cytotoxic killing | - | 10–100 cell units |
| $\theta_i$ | Antigen recognition threshold required for T-cell activation | 5000 | - |
| $\varepsilon$ | Variability in antigen recognition threshold | 0.2 | ±20% |
| $K_{max}$ | Maximum killing capacity of an individual T cell | 5 [#] | - |

Parameters marked by * are estimated from the experimental data in the paper, while parameters marked by [#] are adapted from Bhatt et al[29,30].

### Supplementary Figure 1

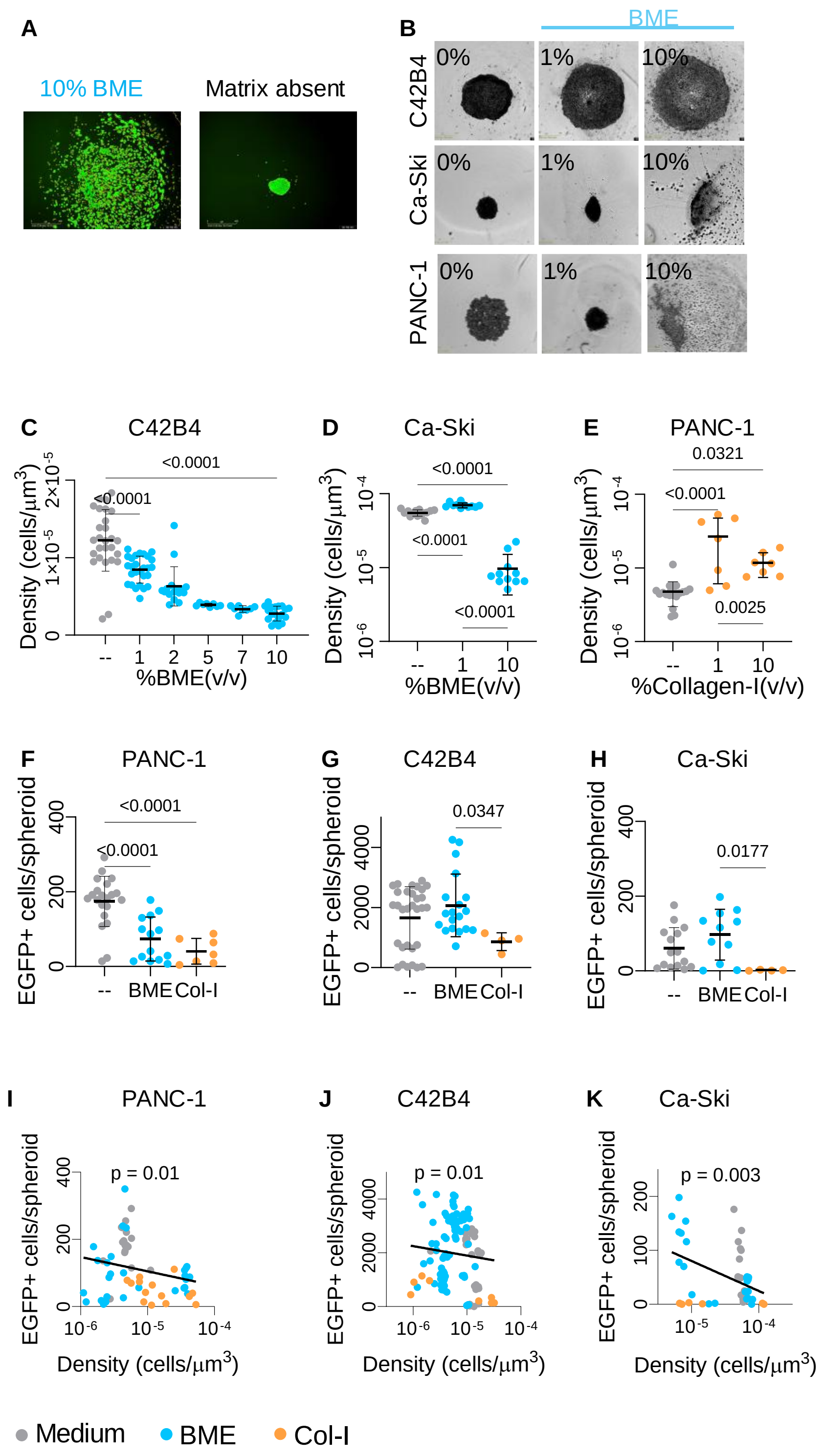


***Figure S1: Effects of extracellular matrix on cell density and infection of different cancer cell types.*** *(**A**) Live-cell images of infected 22Rv1 spheroids when cultured in the presence or absence of extracellular matrix. (**B**) C4-2B4, Ca-Ski, and PANC-1 spheroids captured during culture with 0%, 1%, or 10% BME. (**C–E**) Effect of BME or Collagen-I on spheroid density of different cancer cell lines. (**F–H**) EGFP-positive cells per spheroid comparing no matrix versus 10% BME or 10% Collagen-I for different cell lines. (**I–K**) EGFP-positive cells per spheroid versus cell density after 24 h for different cell lines. Replicates were randomly plated and varied across conditions; all plots used >6 replicates, with some conditions using >9 replicates.*

**Supplementary Figure 2**

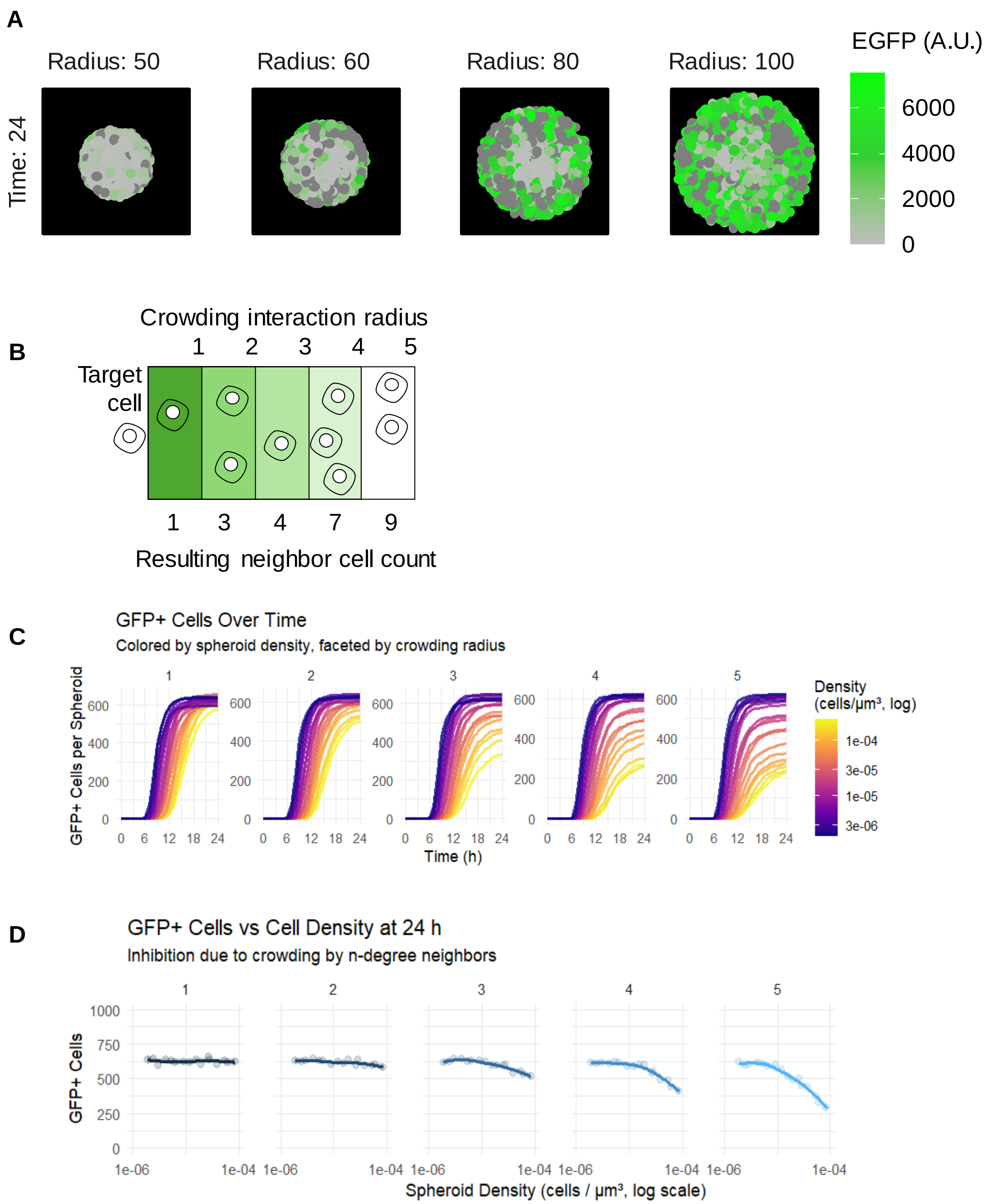


***Figure S2: Computational modeling of infection kinetics.*** *(**A**) shows in silico model snapshots of EGFP expression levels in rSFV-infected cancer cells as a function of spheroid density. (**B**) schematically illustrates how the crowding interaction radius affects the influence neighboring cells exert on EGFP expression in a target cancer cell. For example, a crowding radius of 2 corresponds to 3 neighboring cells, whereas a crowding radius of 5 corresponds to 9 neighboring cells. (**C**) shows the number of EGFP-positive cells per spheroid over increasing density over 24 hours of simulation for different crowding radii. (**D**) shows the frequency of EGFP-positive cells as a function of spheroid density for comparison across crowding radii.*

**Supplementary Figure 3**

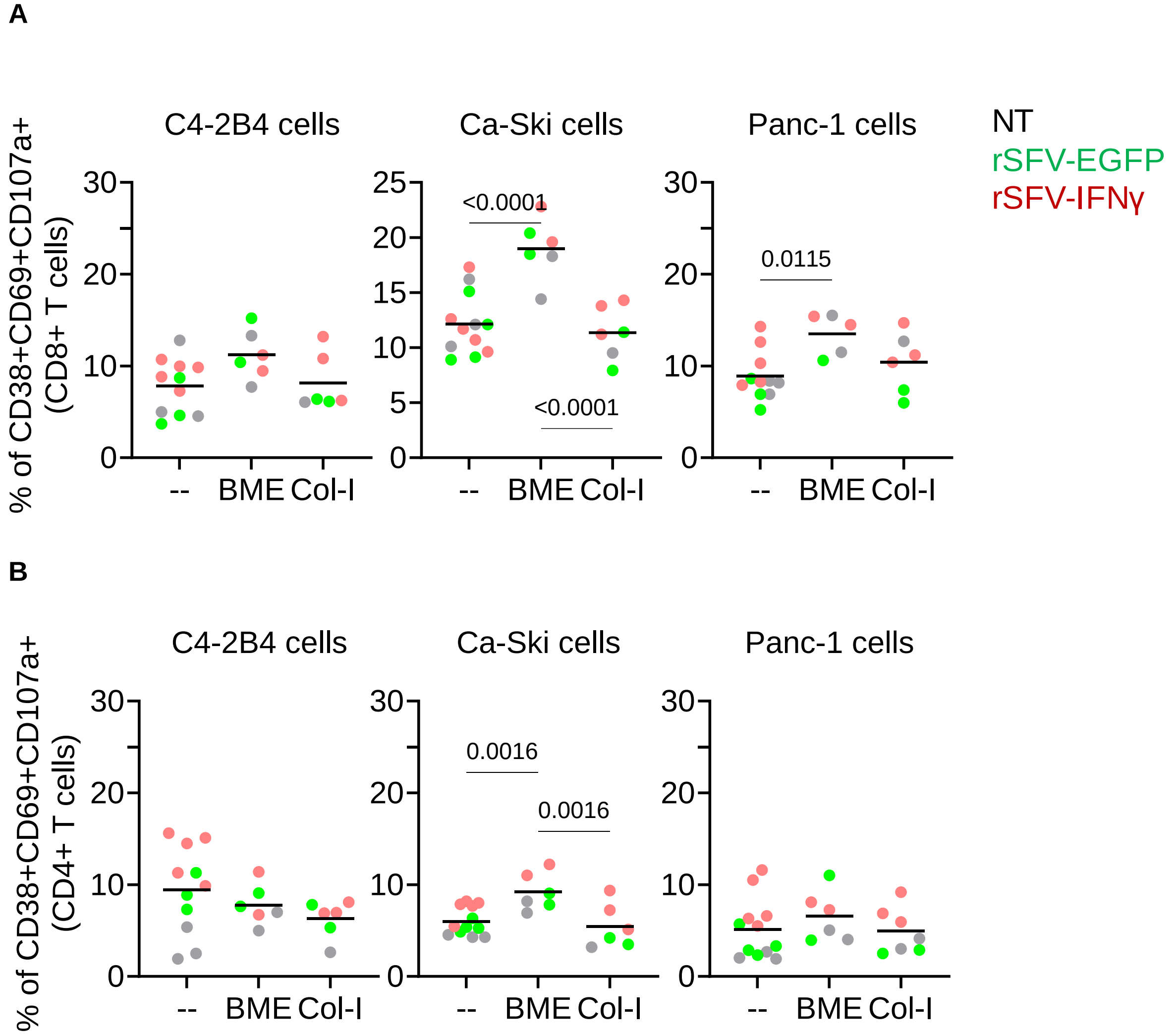


***Figure S3: Effects of extracellular matrix on T cells activation:*** *Activation of (**A**) CD8 and (**B**) CD4 T cells when co-cultured with C4-2B4, Ca-Ski, or PANC-1 cancer cell lines in the presence of BME and Collagen-I extracellular-matrices. Each dot represents a replicate condition (n>6). Colors indicate spheroids status: un-infected (grey), rSFV-EGFP-infected (green) or rSFV-IFNγ-infected (red).*

### Supplementary Figure 4

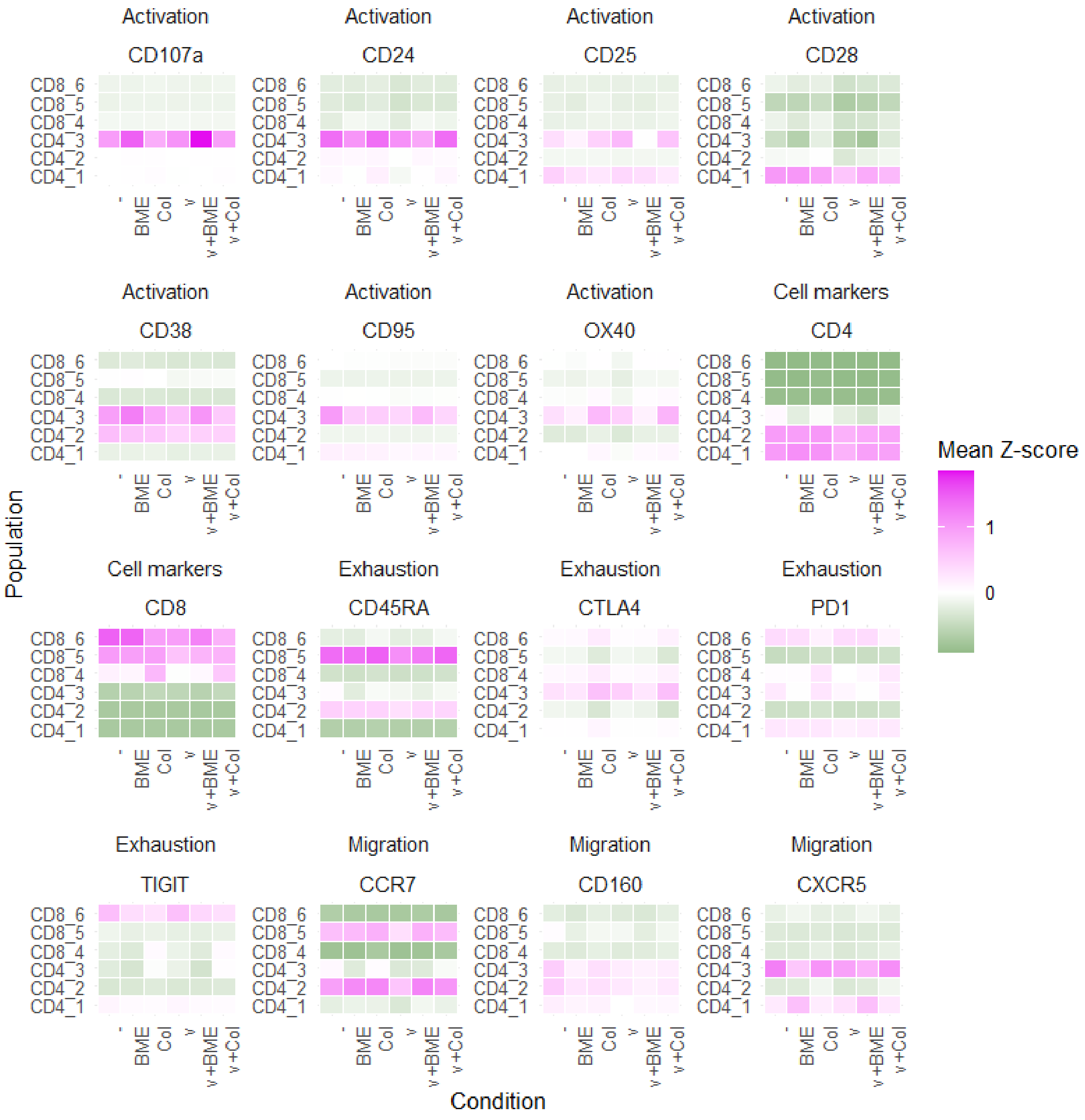


***Figure S4: T-cell phenotype in the presence of extracellular matrix.*** *Surface marker expression (MFI, Z-score) is shown for CD4+ and CD8+ T-cell populations from PBMC–tumor co-culture samples that were either not embedded in matrix or embedded in BME or Collagen-I. Both uninfected (−) and infected (V) samples are included. Surface markers are grouped into three phenotypic categories representing activation, exhaustion, and migration.*

**Supplementary Figure 5**

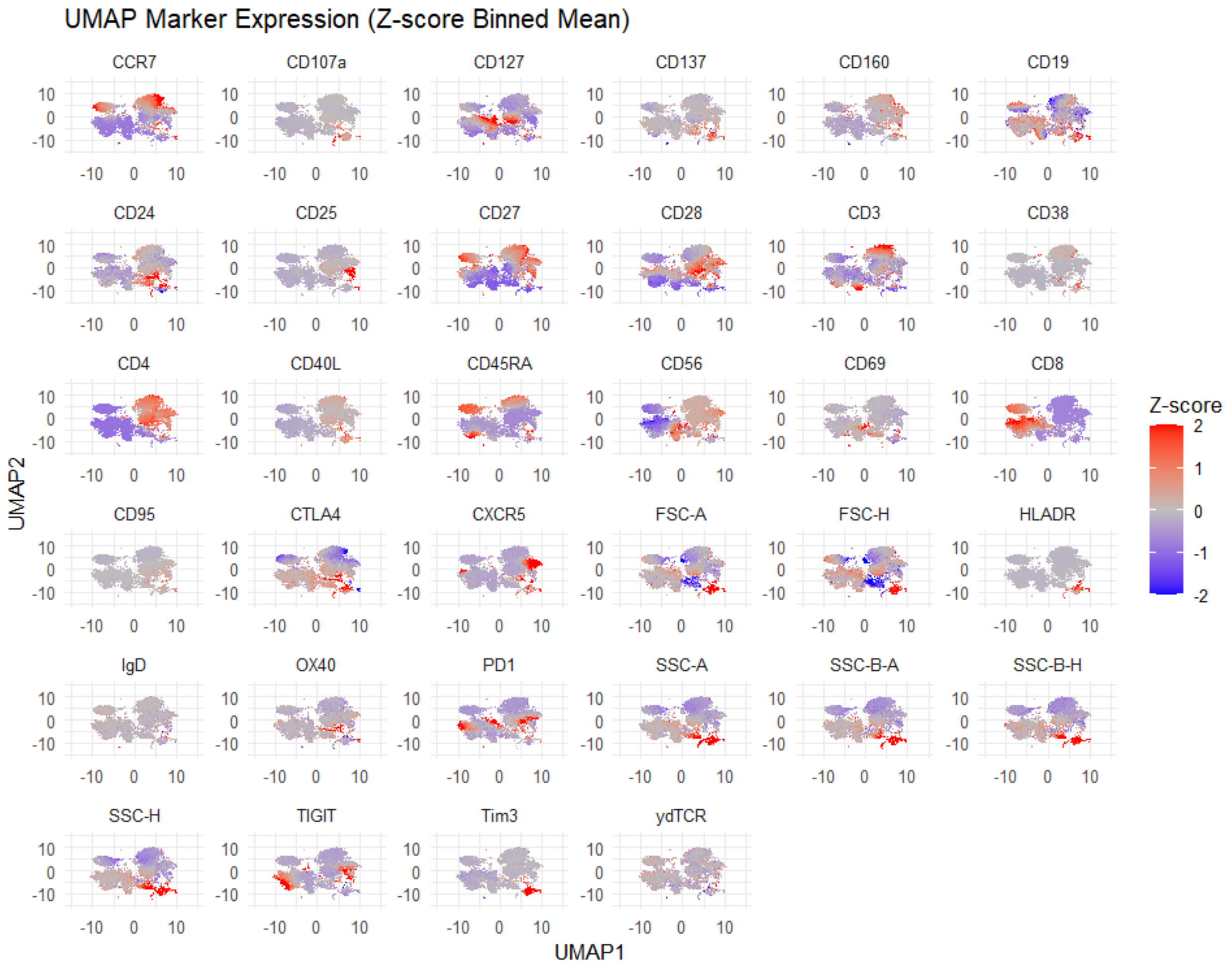


***Figure S5: UMAP marker expression localization.*** *Normalized Z-score of all markers investigated in UMAP cluster of populations. Each marker shown in a separate panel with distribution of cells as points across the different UMAP populations and the color indicating level of marker expression.*
